# GADD45α regulates cell proliferation and DNA repair of BRL-3A cells that treated by FZD/UVC via P38, JNK, CDC2/CCNB1, AKT and MTOR pathways

**DOI:** 10.1101/148759

**Authors:** Xianguang Yang, Lin Zhu, Weiming Zhao, Chuncui He, Shuaihong Li, Cunshuan Xu

## Abstract

GADD45α is a stress-induced gene activated by a variety of stress stimuli, including ultraviolet and ionizing radiation, and involved in cell cycle regulation, apoptosis, maintenance, genomic stability, DNA repair and immune response. However, the effects and regulatory mechanism of GADD45α on proliferation, apoptosis and DNA damage repair of hepatocytes in liver regeneration remains unclear. In this study overexpression of GADD45α significantly inhibited the cell viability, proliferation, the number of cells in G1 and S phases, and of furazolidone (FZD) or UVC induced apoptosis of BRL-3A cells and decreased the inhibition of FZD/UVC on the viability, proliferation of BRL-3A cells, while increased the number of cells in G2/M phase of BRL-3A cells and FZD/UVC induced S phase arrest. Downregulated GADD45α induced the viability, proliferation, the number of cells in S and G2/M phases and the inhibition of FZD/UVC on the viability, proliferation of BRL-3A cells increased, while decreased apoptosis, the number of cells in G1 phases of BRL-3A cells and FZD/UVC induced S phase arrest. The results of qRT-PCR and western blot showed that genes/proteins related to P38MAPK, JNK, CDC2/CCNB1, AKT and MTOR signaling pathways were significantly changed in normal BRL-3A cells. The expression profiles of cell cycle, proliferation and apoptosis related genes/proteins in FZD/UVC treated BRL-3A cells were also detected by qRT-PCR and western blot, and the results indicated that the expression of Myc, Bcl-2, Ccnd1, PCNA, P21, Ccnb1, Caspase3, Caspase8, Caspase9 and Bax have significantly changes.

## 1. Introduction

After partial hepatectomy(PH), the remnant liver cells quickly re-enter the cell cycle from G0 phase, and restore liver quality and function with two cell cycle, which was called liver regeneration (Michalopoulos and DeFrances, 1997). In this process, hepatocytes divided rapidly, which increased the risk of DNA damage during DNA replication (Saintigny et al., 2001). The mechanism of hepatocyte proliferation and DNA damage repair during liver regeneration is not clear.

Growth arrest and DNA damage inducible 45α (GADD45α) is a stress-induced gene and its’ transcription can be activated by various stress, including ultraviolet (Lapeyre et al., 1987), arsenite (Chang et al., 2007), Hypoxia and ionizing radiation, *etc.* and regulate cell cycle (Wang et al., 1999), apoptosis (Gupta et al., 2006), maintain genomic stability, and DNA methylation excision/repair (Li et al., 2015). GADD45α can interact with key cell cycle regulatory factors such as p21 (Kearsey et al., 1995), cdc2/cyclinB1 (Vairapandi et al., 2002), PCNA (Vairapandi et al., 2000) and p38 (Bulavin et al., 2003). The expression of GADD45α in rats liver regeneration(LR) after PH was detected by rat Genome 230 2.0, and the results showed that GADD45α was significantly up-regulated in priming phase (0-6 h) and progressing phase (12-72 h), suggesting that GADD45α may be involved in DNA damage repair and cell proliferation during rat LR after PH.

In this paper, the signal pathway of GADD45α was built and its’ possible mechanism was analyzed by IPA software, QIAGEN and KEGG website. The mechanism of GADD45α on the proliferation, cell cycle, apoptosis and DNA damage repair of BRL-3A cells and BRL-3A cells induced by FZD/UVC were studied by gene overexpression and gene interference. The effects of GADD45α expression on the cell viability, proliferation and apoptosis of BRL-3A and FZD/UVC-induced BRL-3A cells were analyzed by MTT, EdU and flow cytometry. The GADD45α signaling pathways related gene and protein expression profiles were detected by qRT-PCR and western Blot. And the regulatory mechanism of GADD45α on liver cell proliferation, cell cycle, apoptosis and DNA damage repair were analyzed by the above results.

## 2. Materials and methods

### 2.1. Recombinant plasmid construction

TRNzol reagent (TAIGEN) was used to extract total RNA from fresh Rat liver tissue, then the RNA was reverse-transcribed to cDNA with a reverse transcription kit (Promage) for reverse-transcription polymerase chain reaction (RT-PCR). The PCR product of GADD45α (GenBank sequence NM_024127.2, forward primer, 5′- CTCTAGAATGACTTTGGAGG-AATTCTC-3′; reverse primer, 5′-ATAAGAATGCGGCCGCTCACCGTTCGGGGAGATTAA-3′.) was ligated into the pCDH-CMV-MCS-EF1-copGFP Vector. The recombinant plasmid was confirmed by RCR and double restriction enzymes digestion (Not1/XbaI).

### 2.2. Package and harvest of lentivirus that overexpress of GADD45α

The method of Ding et al. (Ding et al., 2016) was used to package the lentivirus. Recombinant plasmid pCDH-CMV-MCS-EF1-copGFP-GADD45α was synthesized by Generay (Shanghai, China). Together with 15 μg helper plasmid pSPAX2, 10 μg helper plasmid pMD2.G, and 50 μL CaCl_2_ (2.5 M), 20 μg recombinant plasmid were added into a clean 1.5 mL microcentrifuge tube, and then sterile water was added to 500 μL, then the above system was added to 500 μL 2×BES (50 mM BES, 280 mM NaCl, 1.5 mM Na_2_HPO_4_) to stand for 20 mins. The medium of HEK293T cells was replaced by DMEM-High glucose medium with 5% serum, and CaCl_2_ system was added into the medium. After 12-16 hours, the medium was replaced with DMEM-High glucose medium with 2% serum. After another 36 hours, the virus particles were collected by passing through a 0.45 μm filter, and were concentrated by ultrafiltration tubes (10,000 MWCO, Millipore, Massachusetts, USA) at 4000 g (4°C) for 20 mins. The concentrated virus particles were suspended in PBS and stored at -80°C.

### 2.3. Transduction of BRL-3A cell

Transduction was performed in 24-well plates. BRL-3A cells were seeded at 5×10^4^ cells per well. One day later, the cells were transduced with 2×10^5^ TU virus particles. 24 hours post transduction, the medium was replaced with DMEM-High glucose medium with 10% serum. After another two days, the transduction efficiency was measured by detecting the green fluorescent protein (GFP) under fluorescence microscope. BRL-3A cells transfected with recombinant plasmid pCDH-CMV-MCS-EF1-copGFP-GADD45α (experimental group, EG) and empty vector pCDH-CMV-MCS-EF1-copGFP (Normal control, NC) were collected after culturing in large scale. qRT-PCR and western blot were used to detect the expression changes of genes/proteins of GADD45α during EG and NC groups.

### 2.4. siRNA transfection of Cells

BRL-3A cells were seeded at 5×10^4^ cells per well. The next day when the cells were 60–70% confluent, the culture medium was replaced by fresh DMEM. 3μL Lipofectamine^®^ RNAiMAX Reagent and 50 pM siRNA were added into 50 μL DMEM respectively. Then Add diluted siRNA to diluted Lipofectamine^®^ RNAiMAX Reagent (1:1 ratio) and incubate for 5 minutes at room temperature. Finally, 50 μL siRNA-lipid complex were added into one well which contained 450 50 μL DMEM. Then the cells were cultured for 6 hours, and its’ media were changed to fresh complete DMEM. The cells were harvested at 48 h after transfection.

### 2.5. FZD treatment and UV irradiation

FZD was dissolved in dimethyl sulfoxide (DMSO) to make a stock solution of 50 mg/mL and further diluted to corresponding concentrations with the cell culture medium. BRL-3A cells were cultured in the medium as mentioned above in 96-well plates at a density of 1.5×10^4^ cells per well. After culture for 24 h, the cells were exposed to different concentrations of FZD (0, 12.5, 25,50 and 100 μg/mL) or UV light is 30cm below the UVC irradiation 0, 15, 30, 45 and 60 s. The cell viability was determined by MTT assay as previously described. The absorbance at 490 nm at 24 h after treatment was measured by Biotek reader (ELx800, USA). The cell viability and cytotoxicity were estimated as the percentage of the control

### 2.6. Comet Assay

DNA damage was evaluated through the alkaline single cell gel electrophoresis (comet) assay. 5×10^6^ cells were collected and added to 100 PBS. 100 μL 0.8% normal elting point agarose was dropped onto slides, and t covered with a glass overslip. The slides were maintained at 4 °C for 10 min; verslips were removed, and 15 μL of treated cells were mixed with 60 μL of 0.6% low melting point agarose in PBS at 37 °C and spread onto slides for 10 min at 4 °C. The final ayer of 70 μL of 0.6% low melting point agarose was applied in the same way. The slideswithout coverslips were immersed in a cell lysate with 10% DMSO at 4 °C for 2 h and then placed in electrophoresis buffer (1 mmol/L of EDTA, 300 mmol/L of NaOH, pH > 13) in an electrophoresis tank for 20 min to allow alkaline unwinding. Electrophoresis was performed for 20 min under 25 V and 300 mA. The slides were then transferred to 0.4 mmol Tris-buffer (pH 7.5), washed twice for 10 mins. Comets were stained with propidium iodide (2 μg/mL) and analyzed under a confocal laser scanning microscope (Nikon, Tokyo, Japan). Image analysis and tail moment were performed with CASP 1.2.2 software (Wroclaw, Silesia, Poland); 25 cells were randomly selected per sample.

### 2.7. Cell Proliferation Assay

Cells were treated by the methods above. Then 10 μL MTT solution (5 mg/mL in PBS) (Geneview, USA) were added to each well at 24, 48 and 72 h, and the cells were incubated for 4 hours in the dark at 37 °C. Then, the medium was replaced with 100 μL dimethylsulfoxide (DMSO) (Sigma-Aldrich, USA) for 10 mins to dissolve the formazan crystals produced by cells. Subsequently, the absorbance at 490 nm was measured by Biotek reader (ELx800, USA).

EdU detection was performed as the 5-ethynyl-2’-deoxyuridine (EdU) labeling/detection kit (Ribobio, Guangzhou, China). Cells were seeded and treated with the methods above, after 46 hours, the medium were replaced with new DMEM-High glucose medium containing 50 μM EdU for 2hr incubation at 37°C under 5% CO_2_. After fixation with 4% paraformaldehyde (pH 7.4) for 30 min and glycine for 5 min, anti-EdU working solution was added for reaction at room temperature in dark for 30 mins. Then cells were washed with 0.5% TritonX-100 and incubated with 5 g/ml Hoechst 33342 dye at room temperature in dark for 30 mins. Cells were visualized under a fluorescence microscope.

### 2.8. Flow cytometric analysis

Cells were collected at 48 h after treatment, and washed with PBS followed by fixation in 70% ethanol for 12 hours on ice. After washing with PBS, cells were incubated in PBS with 0.2 mg/mL RNase (Solarbio, China) at 37 °C for 30 mins. Then cells were stained with PI (20 μg/ml, Sigma, USA) in dark for 15 mins. Finally, the stained nuclei were counted by flow cytometry and the cell cycle profile was analyzed by the FlowJo software.

Cells were subsequently harvested with 0.25% trypsin without EDTA, washed twice with cold PBS and resuspended in 500 μL binding buffer(1 × 10^6^ cells/mL). Then cells were incubated with 5 μL PE Annexin V (40 μg/mL) and 5 μL 7-AAD (40 μg/mL) in the dark for 15 min at room temperature and the cell apoptosis condition were detected by flow cytometry.

### 2.9. Identification of GADD45α signaling pathway

Previous studies have reported that GADD45α could be activated by UV/Ionizing Radiation, and then exert its function via CDC2/CCNB1, mTOR, P21, and PCNA signaling pathway. Therefore, “CDC2/CCNB1 signaling pathway”, “mTOR signaling pathway”, “P21 signaling pathway”, “PCNA signaling pathway” and were entered into the “Pathway and Function” of Ingenuity Pathway Analysis (IPA) software respectively to obtain the interaction network of them and GADD45α, and the above results were filtered and integrated by researching the signaling pathway maps at the databases including GenMAPP, KEGG, BIOCARTA, QIAGEN and Biocompare, and then network of GADD45α-mediated signaling pathway were reconfirmed by pertinent articles.

### 2.10. qRT-PCR

Cells were seeded and treated according to the methods above and collected at 48 h after the treatment. The total RNA was extracted using TRNzol according to the manufacturer’s instructions. Total RNA (2 μg) was reverse-transcribed using random primers and reverse transcription kit (Promega, Madison, USA).The primers for the genes involved in GADD45α signaling pathway were designed by Primer Express 5.0 software (**Tab 1**). The PCR was performed under the conditions with SYBG I: 2 min at 95 °C, followed with 40 cycles for 15 s at 95 °C, 15 s at the corresponding annealing temperature, and 30 s at 72 °C. *β-actin* was used as an internal reference. Each sample was performed in triplicates.

**Tab 1.**
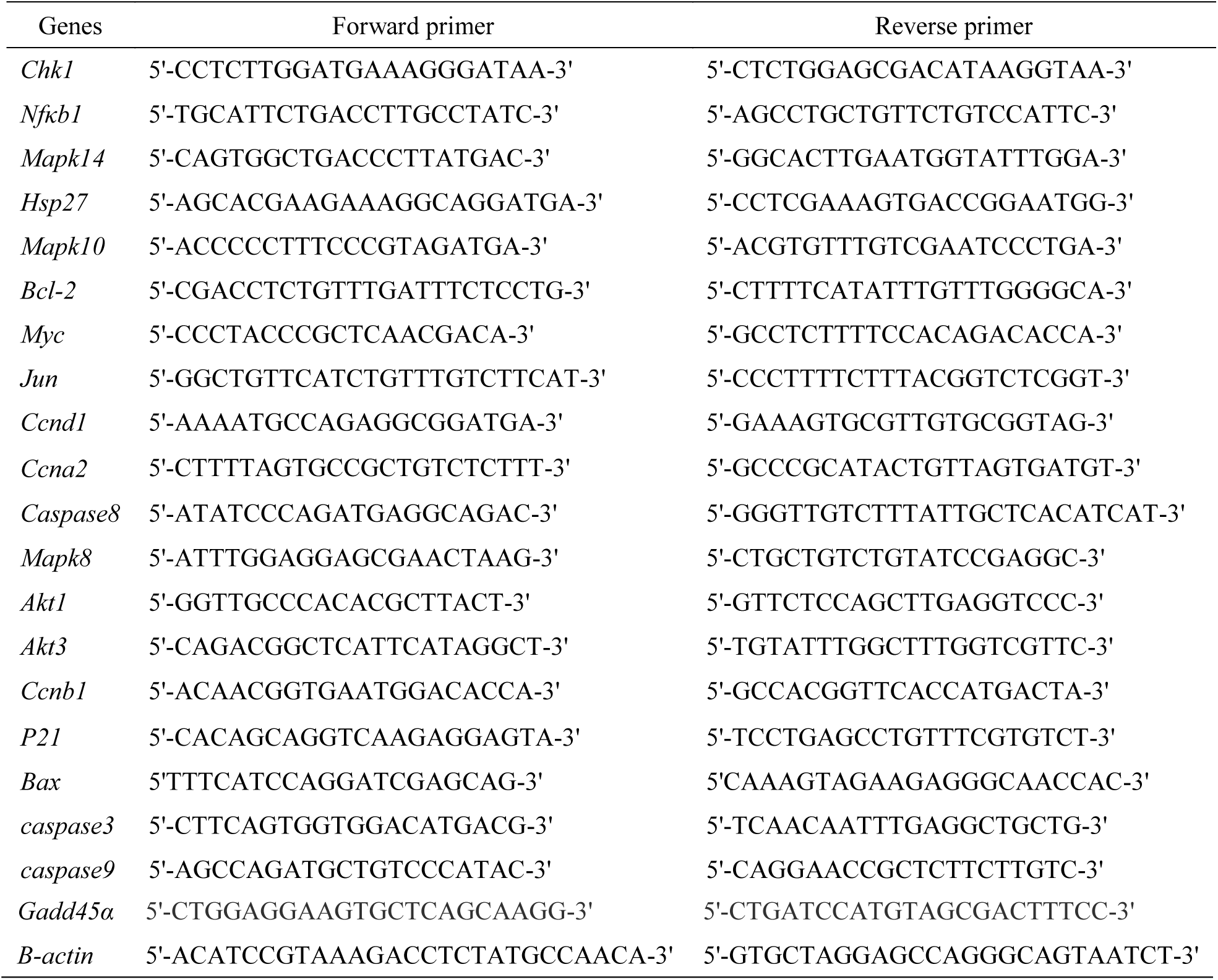
The primer sequences of genes for qRT-PCR

### 2.11. Western blot analysis

Cells were seeded and collected as described in the above methods, equal amounts of total proteins (20μg) were separated by SDS-PAGE, and then transferred onto a nitrocellulose membrane(“PALL Corporation”, USA).After the transfer, the membrane was blocked with 5% skimmed milk in Tris-buffered saline (TBS) for 1 hour and subsequently incubated overnight at 4°C with the antibodies (β-ACTIN, NF-ΚB1, JUN, CCNB1, BCL-2, MYC, CASPASE3, CASPASE8, CASPASE9, PCNA and P21, Cell Signaling Technology) against proteins-related to GADD45Α signaling pathway at a dilution of 1:500. After washing with TBS containing 0.02% Tween-20 for 30 min at room temperature, the membrane was further incubated with alkaline phosphatase labelled secondary antibodies goat anti-rabbit/mouse IgG (“Sigma,” Germay) at a dilution of 1:1 000 for 1 hour. Finally, protein bands were visualized with BCIP/NBT. Image QuantTMTL (“Sigma”, Germay) was used to measure the relative abundances of target proteins. β-ACTIN at a dilution of 1:1000 was served as an internal reference.

### 2.12. Statistical analysis

All the experiments were repeated for three times. Data was expressed as mean ± SD. One-way analysis of variance (ANOVA) followed by a LSD test was used to compare experimental values between groups. Symbol * indicates *P*<0.05 and symbol ** *P*<0.01.

## 3. Results

### 3.1. Identification of GADD45α overexpression vector

The recombinant plasmid was confirmed by RCR and double restriction enzymes digestion, and the results showed that the length of PCR and double restriction enzymes digestion product were consistent with the known GADD45α gene (Fig 1).

**Fig 1.**
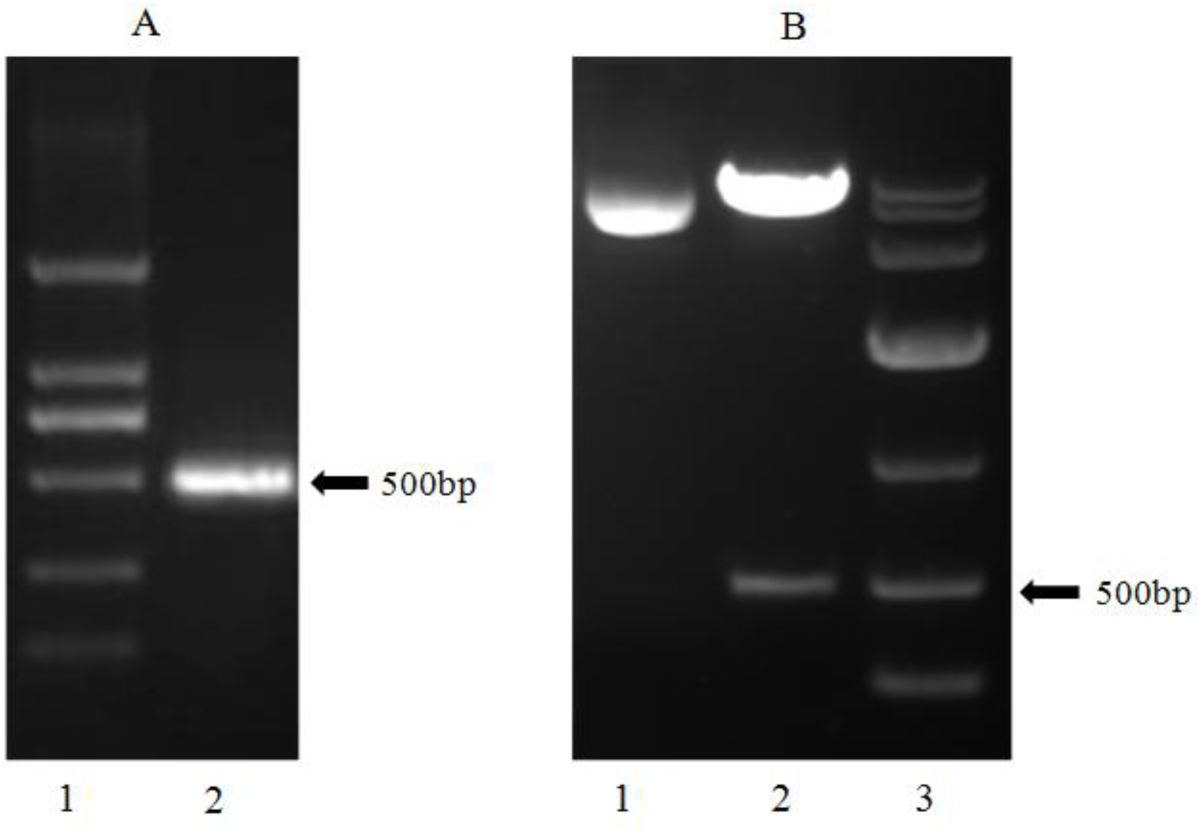
Identification of GADD45α recombinant plasmid. A: PCR identification, 1: 2000 Marker, 2: Purpose band, B: Identification of double restriction enzymes digestion, 1: PCDH-CMV-MCS-EF1-copGFP-GADD45α. 2: Not1/XbaI double restriction enzymes digested, 3: 10000 Marker.

### 3.2. GADD45α was highly expressed in BRL-3A cells

The transduction efficiency was measured by observance of the GFP under fluorescence microscope at 48 h after the transduction. The results showed that recombinant plasmid pCDH-CMV-MCS-EF1-copGFP-GADD45α was transduced into the BRL-3A cells successfully (Fig 2). qRT-PCR and Western blot were used to detect the expression of GADD45α in EG and NC groups, and the results indicated that the expression of GADD45α in EG was markedly increased than that of NC(Fig 3).

**Fig 2.**
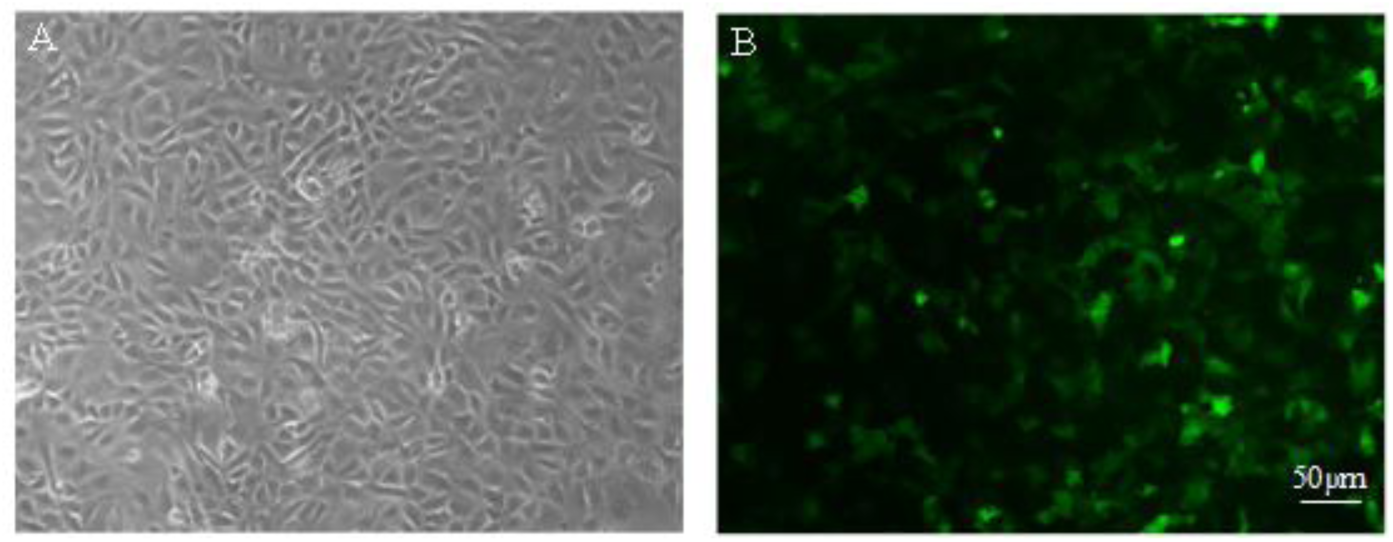
Efficient infection with recombinant plasmid in BRL-3A cells. A: white light, B: green fluorescence

**Fig 3.**
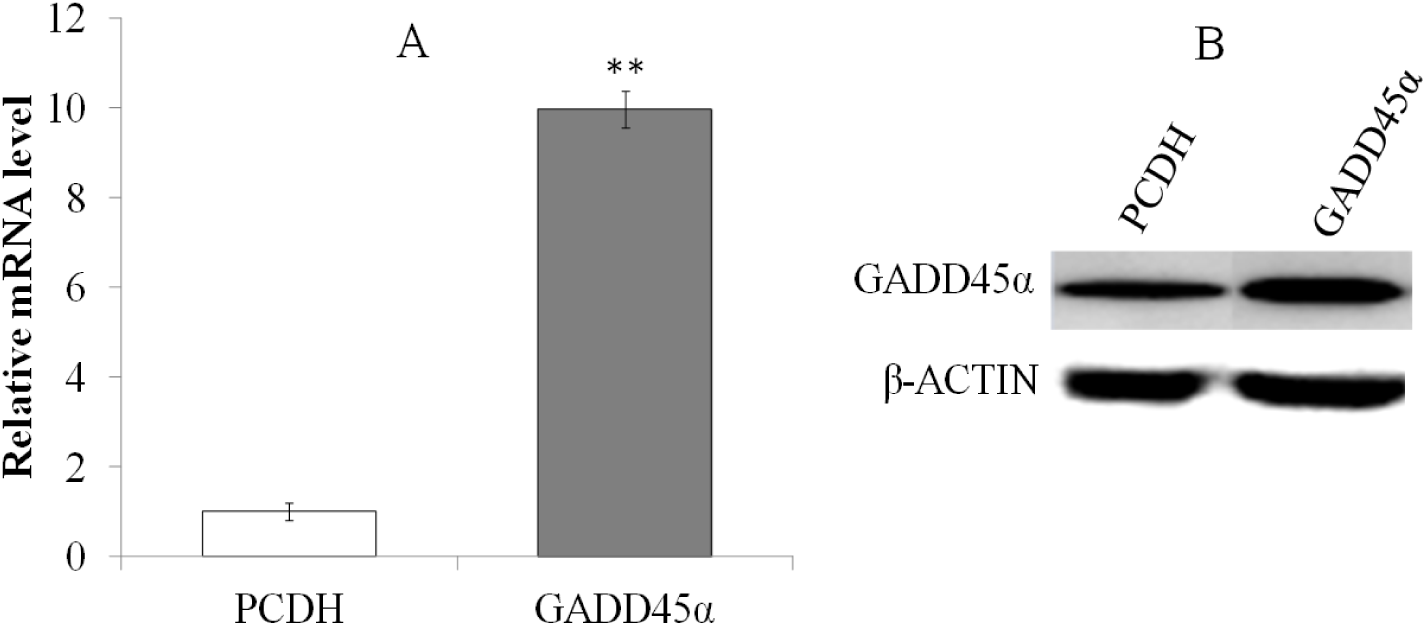
Expression changes of GADD45α between EG and NC. A: qRT-PCR, B: Western blot. \*\**p*<0.01

### 3.3. Suppression of GADD45α expression by siRNA in BRL-3A Cells

The Negative control (NC)、 siRNA1、 siRNA2、 siRNA3 was transfected into BRL-3A cells, and the interference efficiency on GADD45α mRNA levels were determined by comparison with control group by quantitative real-time PCR at the time points, and the results indicated that their interference efficiency were 56.47±2.31%, 12.95±2.65%, 24.21±4.58%(Fig. 4). Accordingly, siRNA1 was selected as an effective interference fragment for the later study.

**Fig 4.**
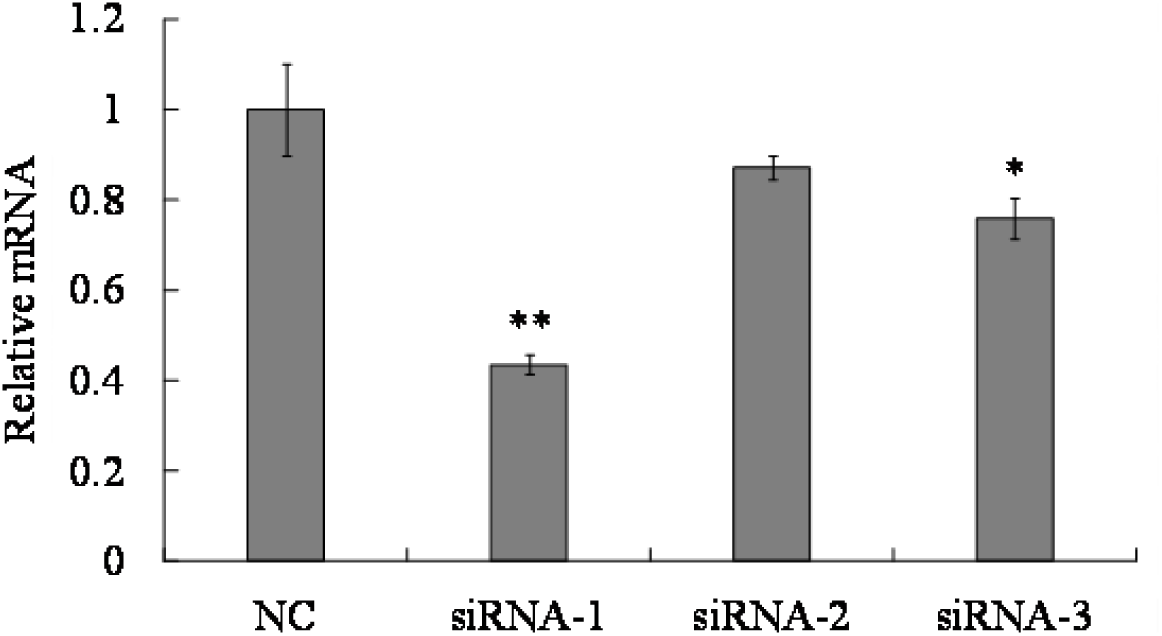
The interference efficiency of siRNAs. \**p*<0.05; \*\**p*<0.01

### 3.4. Cell viability inhibition by FZD or UVC

After treating with 12.5, 25, 50 and 100 μg/mL FZD, cell viability at 24 h were decreased by 22.12±1.02%, 37.15±2.12%, 49.91±2.51% and 77.94±1.34%, respectively. Cells were exposed to UVC for 15, 30, 45 and 60 s, and cell viability decreased by 31.42±0.95%, 49.13±2.01%, 75.76±2.32% and 80.52±1.57% at 24 h after treatment respectively. Accordingly, 50 μg/mL FZD and exposed to UVC for 30 s were optimized to be as an effective condition for the later study.

**Fig 5.**
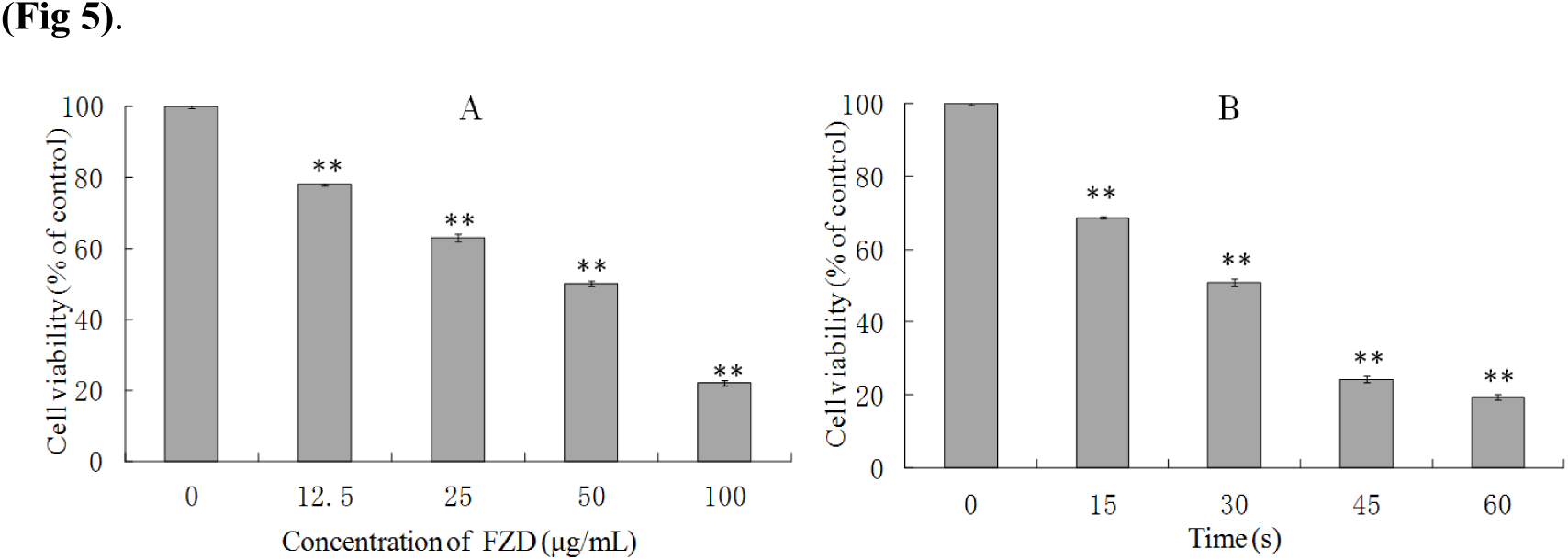
The survival ratios of BRL-3A cells after FZD (A) and UVC (B) treatment. \*\**p*< 0.01, compared with control group.

### 3.5. FZD and UV-Induced DNA Damage by Comet Assay

DNA fragment that caused by FZD treatment and UV irradiation was demonstrated by comet assay (Fig 6). The fragmented DNA, especially the small fragments of DNA, can enter agarose gel easily, the more (smaller) fragments, the longer the tail becomes. The results indicated that the nuclear of the control group were round and there were no tailing, and there were obvious tailing in the experiment groups treated with FZD and UVC irradiation. The results showed that both FZD treatment and UVC irradiation will lead to DNA damage.

**Fig 6.**
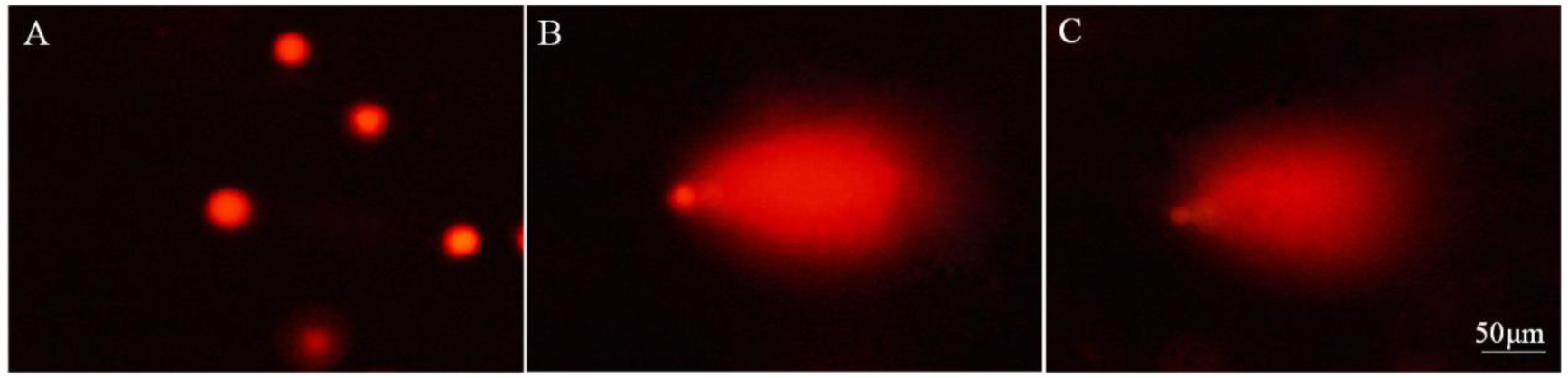
FZD and UVC-induced DNA damage detected by comet assay. A: Control, B: FZD treatment, C: UVC-irradiation.

### 3.6. GADD45α effect cell proliferation of BRL-3A and FZD/UVC-stimulated cell growth

#### suppression

EdU and MTT were used to test the role of GADD45α on BRL-3A cell proliferation and FZD/UVC-stimulated cell growth suppression. Result of EdU assay showed that GADD45α overexpression inhibited BRL-3A cell proliferation and attenuated the inhibition of FZD/UVC-induced proliferation of BRL-3A cells, whereas downregulation of GADD45α promotes cell proliferation and enhances the inhibition of FZD/UVC-induced proliferation of BRL-3A cells (Fig 7).

**Fig 7.**
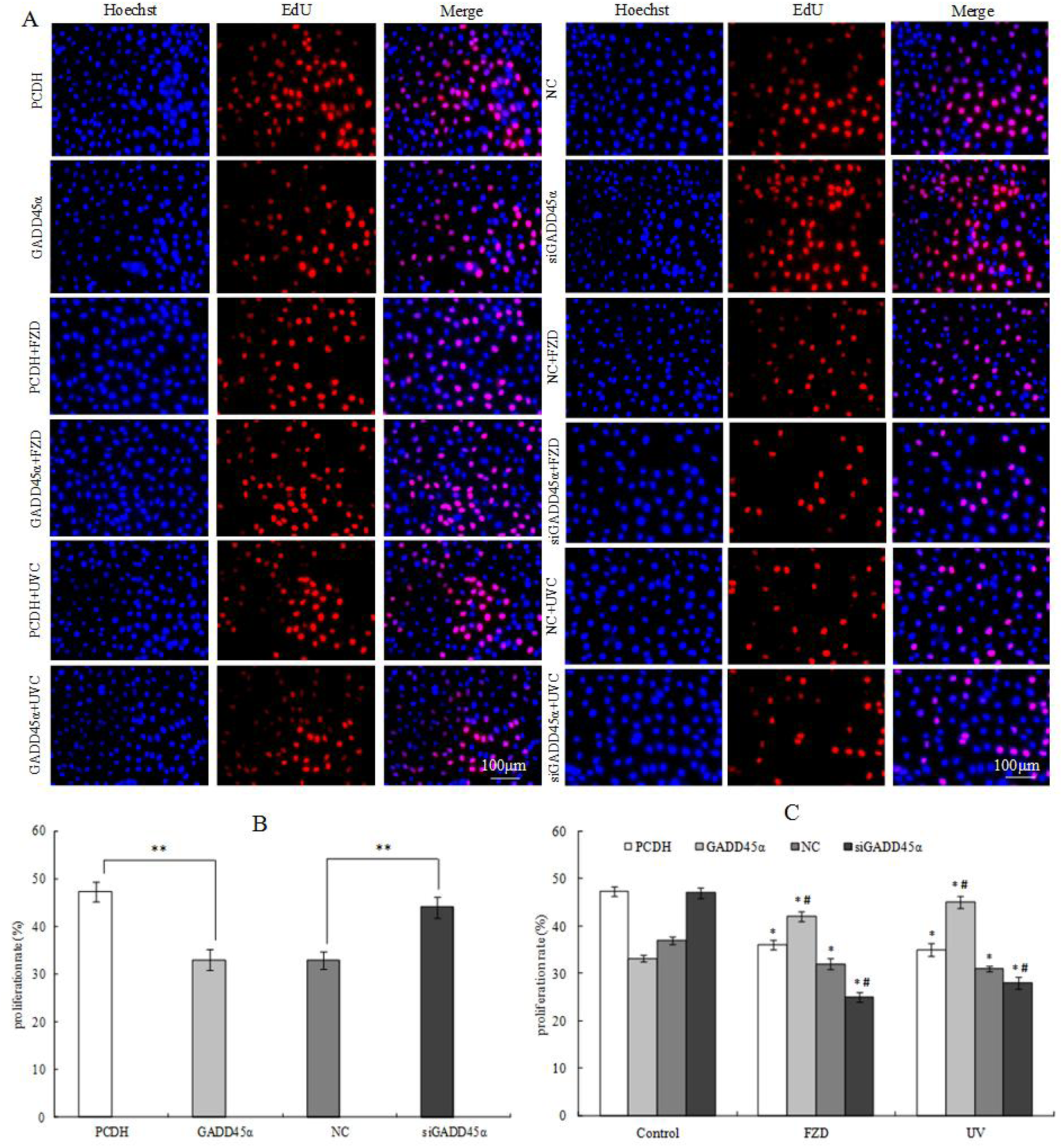
Effects of GADD45α expression changes on cell proliferation of BRL-3A cells and cells treated by FZD/UVC A: Effects of GADD45α expression changes on cell proliferation of BRL-3A cells and cells treated by FZD/UVC; B: Effects of GADD45α expression changes on cell proliferation of BRL-3A cells; C: Effect of GADD45α expression changes on cell viability of FZD and UVC-treated cells. * *p*<0.05, \*\**p*<0.01 compared with control; #*p*<0.05 compared with control + FZD / UVC

MTT assay result showed that the up-regulation of GADD45α gene inhibited the cell viability of BRL-3A cells and promoted viability of BRL-3A that treated by FZD/UVC. While down-regulation of this gene increased the cell viability of BRL-3A cells and reduced the viability of BRL-3A cells treated by FZD and UVC (Fig 8).

**Fig 8.**
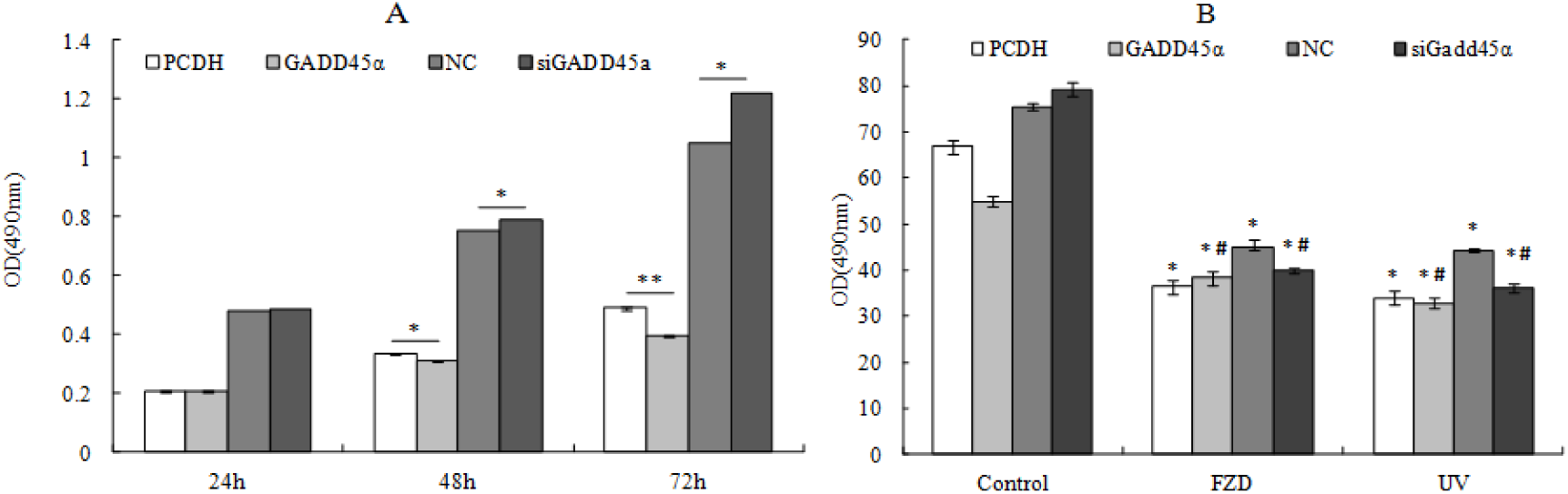
Effects of GADD45α expression changes on cell viability of BRL-3A cells and BRL-3A cells treated by FZD and UVC A: Effects of GADD45α expression changes on cell viability of BRL-3A cells B: Effect of GADD45α expression changes on cell viability of FZD and UVC-treated cells * *p*<0.05; \*\**p*<0.01 compared with control; #*p*<0.05 compared with control + FZD/UVC

### 3.7. The effect of GADD45α on cell cycle distribution

GADD45α may affect FZD-stimulated S phase cell cycle arrest through a signal transduction network. To test the role of GADD45α in BRL-3A cells cell cycle and FZD or UVC-stimulated S phase cell cycle arrest, BRL-3A cells were exposed to 50 g/mL FZD for 24 h and UVC for 30s, cell cycle were monitored by flow cytometry. As shown in Fig. 9 FZD and UVC could obviously arouse S-phase cell cycle arrest in BRL-3A cells. When GADD45α low-expressing BRL-3A cells were exposed to FZD for 24 h and UVC for 30s, proportion of cells in S phase decreased from 43.7±0.70% and 37±0.38% to 31.5±0.50% and 27.5±0.42% respectively (*p* < 0.05), and G2/M phase increased from 5.96±0.45% and 11±0.53% to 11.2±0.45% and 18.3±0.38% (P < 0.05), whereas G1 phase increased from 50.3±0.60% and 52±0.30% to 57.4±0.51% and 54.2± 0.40%, compared to that of control cells induced by FZD and UVC (Fig. 9, 10). These results indicated that GADD45α could modulate FZD/UVC-induced S-phase cell cycle arrest.

**Fig 9.**
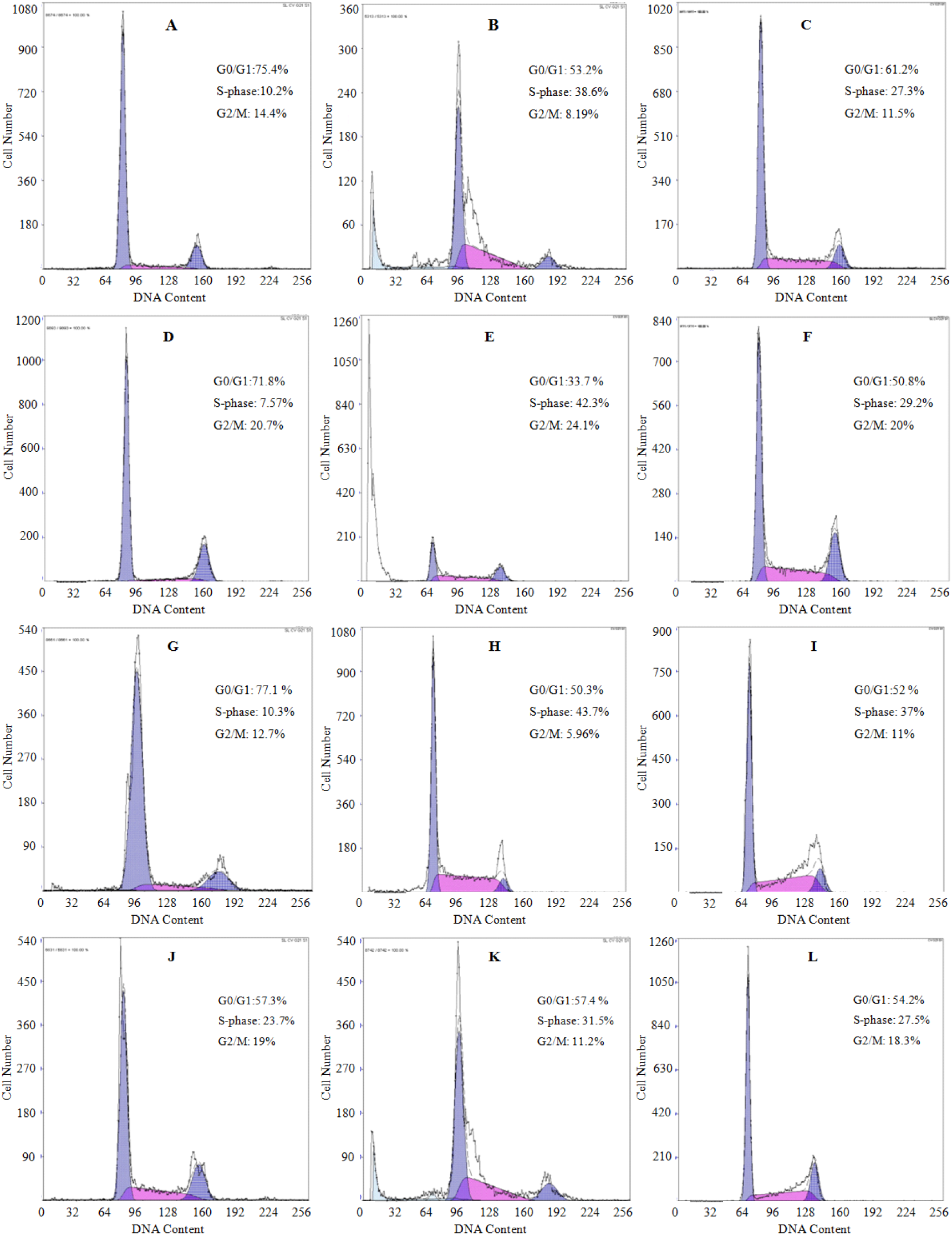
Effects of GADD45α expression changes on cell cycle of BRL-3A cells and BRL-3A cells treated by FZD /UVC. A: PCDH, B: PCDH+FZD, C: PCDH+UVC, D: GADD45α, E: GADD45α+FZD, F: GADD45α+UVC, G: NC, H: NC+FZD, I: NC+UVC, J: siGADD45α, K: siGADD45α+FZD, L: siGADD45α+UVC.

**Fig 10.**
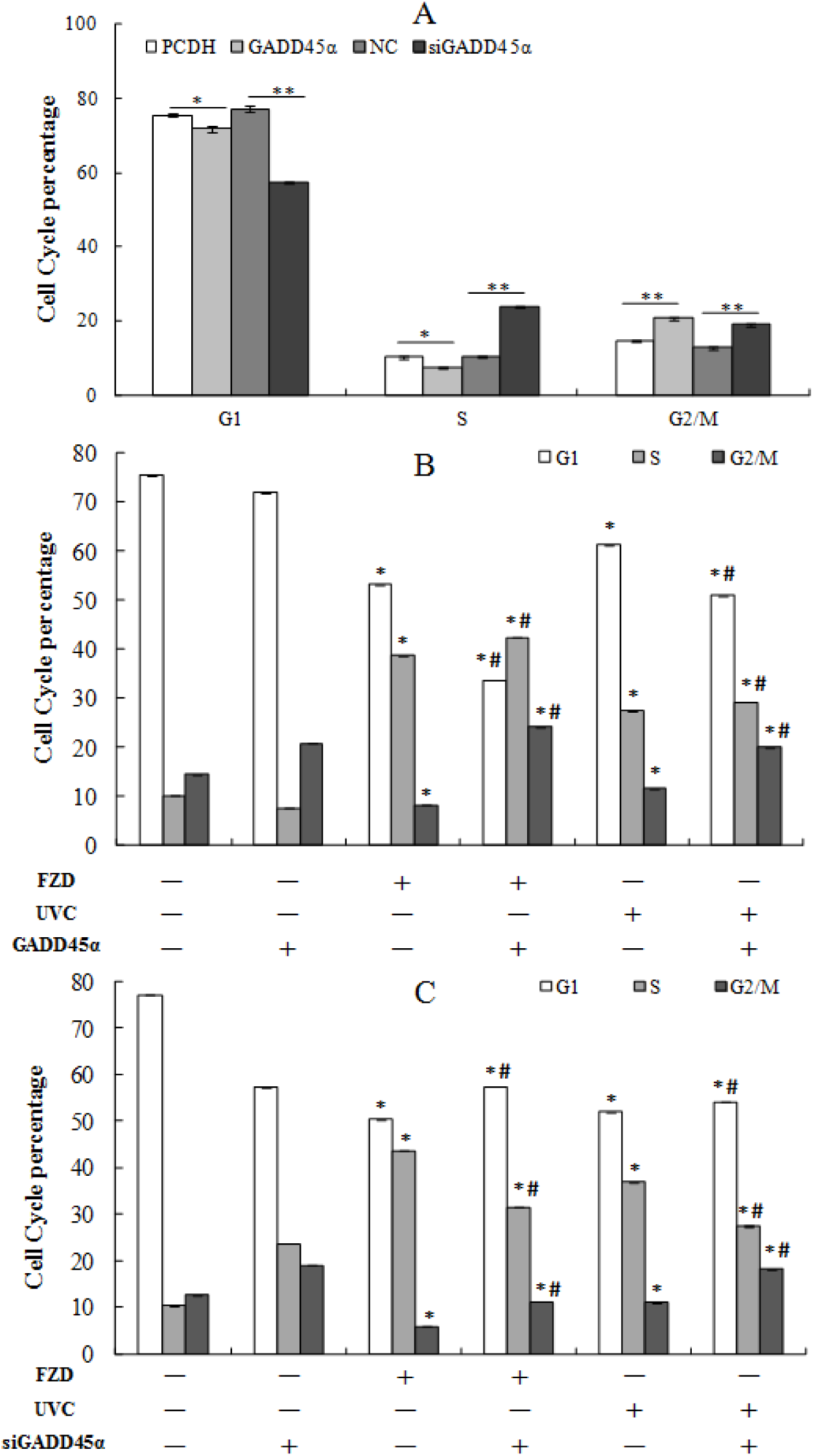
Analysis of the effect of GADD45α expression on cell cycle of BRL-3A cells and BRL-3A cells treated by FZD/UVC. A: Effect of GADD45α expression change on cell cycle; B: Effect of GADD45α expression up-regulation on cell cycle arrest induced by FZD and UVC; C: Effect of GADD45α down-regulation on FZD and UVC-induced cell cycle arrest. * *p*<0.05 compared with control; #*p*<0.05 compared with control + FZD/UVC.

The results showed that overexpression of GADD45α significantly inhibited the number of cells in G1 and S phases, while increased the number of cells in G2/M phase of BRL-3A cells and FZD/UVC induced S phase arrest. Downregulated GADD45α induced the number of cells in S and G2/M phases, while decreased the number of cells in G1 phases of BRL-3A cells and FZD/UVC induced S phase arrest.

### 3.8. Effect of GADD45α expression change on cell apoptosis

The results showed that overexpression of GADD45α have no effect on the apoptosis of BRL-3A cells, while significantly inhibited the cell apoptosis that induced by FZD/UVC. Downexpression of GADD45α significantly inhibited the apoptosis of BRL-3A cells, while significantly promoted the cell apoptosis that induced by FZD/UVC (Fig. 11, 12).

**Fig 11.**
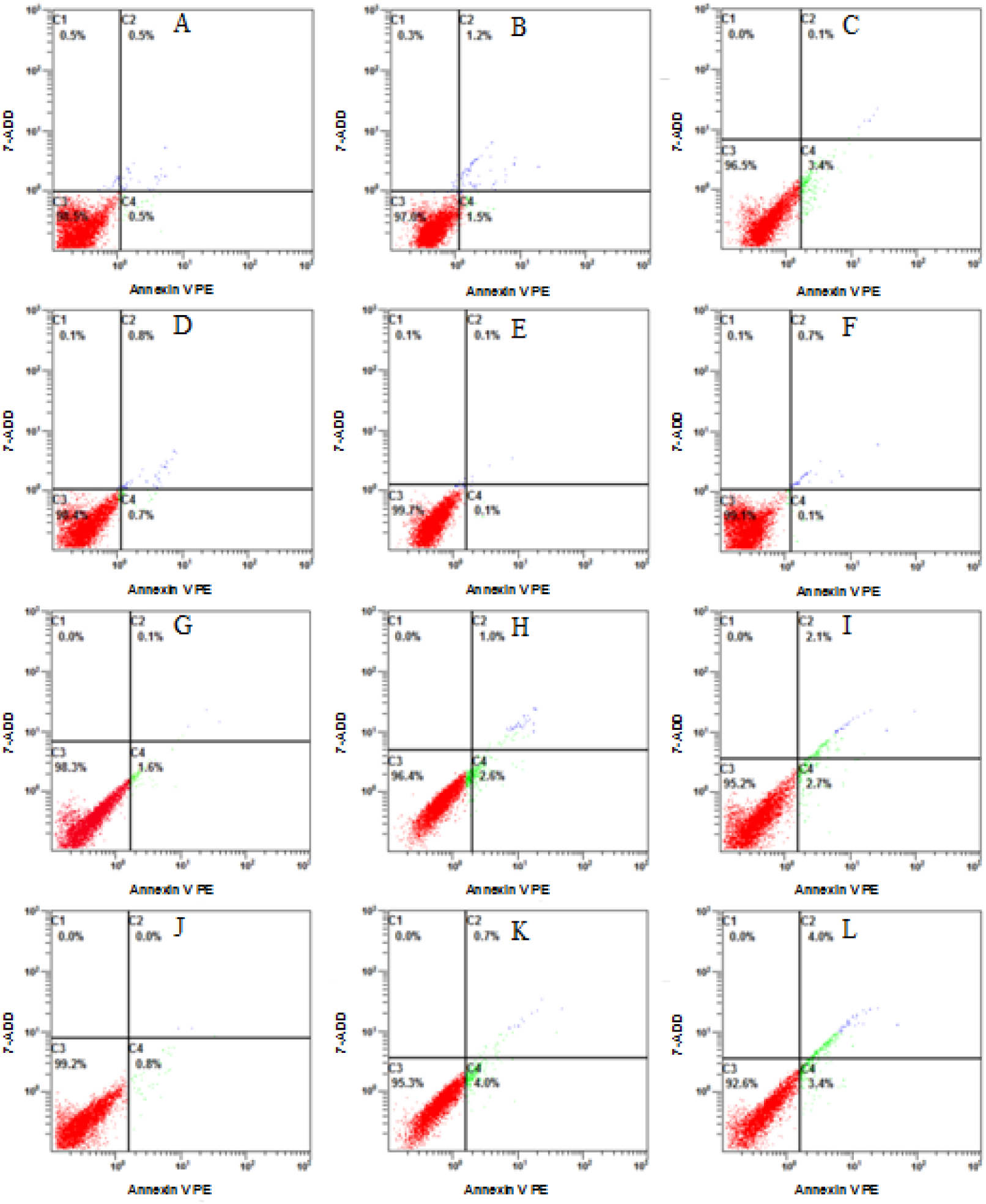
Effects of GADD45α expression change on the apoptosis of BRL-3A cell and FZD/UV-induced apoptosis A: PCDH, B: PCDH+FZD, C: PCDH+UVC, D: GADD45α, E: GADD45α+FZD, F: GADD45α+UVC, G: NC, H: NC+FZD, I: NC+UVC, J: siGADD45α, K: siGADD45α+FZD, L: siGADD45α+UVC.

**Fig 12.**
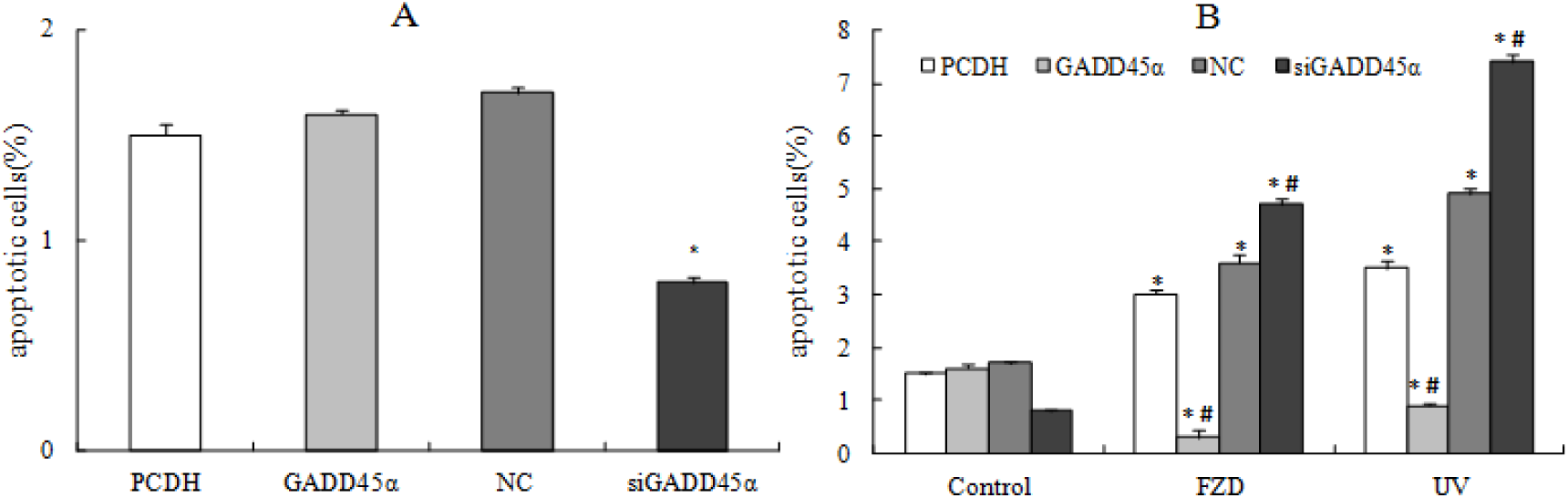
**Apoptosis** percentage of treated cells. A: Effect of GADD45α expression change on BRL-3A cells, B: Effect of GADD45α expression change on apoptosis of BRL-3A cell that treated by FZD/UVC. * *p*<0.05 compared with control; #*p*<0.05 compared with control + FZD/UVC

### 3.9. GADD45α signaling pathway

GADD45α signaling pathway was built by the IPA software, and the results showed that GADD45α could regulate proliferation of cells, DNA repair, G2/M phase, and apoptosis response through P38MAPK, JNK, CDC2/CCNB1, AKT and MTOR signaling pathways (Fig 13).

**Fig 13.**
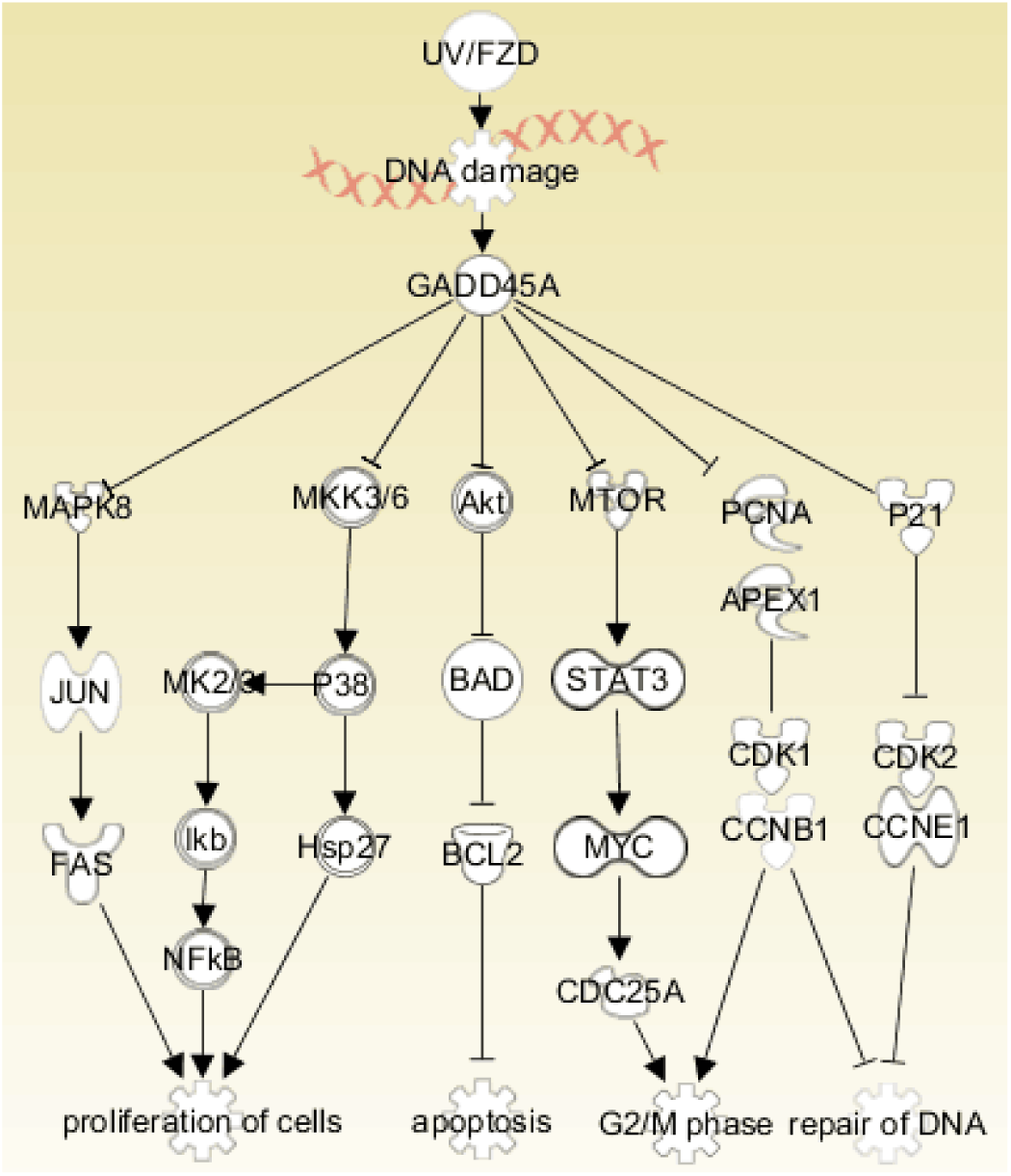
GADD45α signaling pathway.

### 3.10. GADD45α expression changes affected the expression of genes/proteins related to **GADD45α signaling pathway**

To further discover the underlying mechanism of GADD45α in normal BRL-3A cells and FZD/UVC-induced BRL-3A cells on cell cycle and apoptosis, the expression changes of genes-associated with GADD45α pathway were detected. The results indicated that overexpression of GADD45α in BRL-3A cells increased the expression of Mapk8, Mapk10, Mapk14, Hsp27, Akt1/3, Myc, Bcl2, NF-κB1, Ccna2 and Ccnd1 were down-regulated at mRNA and protein, while the expression of apoptosis gene-related gene Caspase8 was up-regulated. When down-regulated the expression of GADD45α, the expression of Mapk8/10/14, Hsp27, Akt1/3, Myc, Bcl-2, NF-κB1, Ccnd1 and Ccna2 were up-regulated at mRNA and protein, while the expression of Caspase8 was down-regulated (Fig 14).

**Fig 14.**
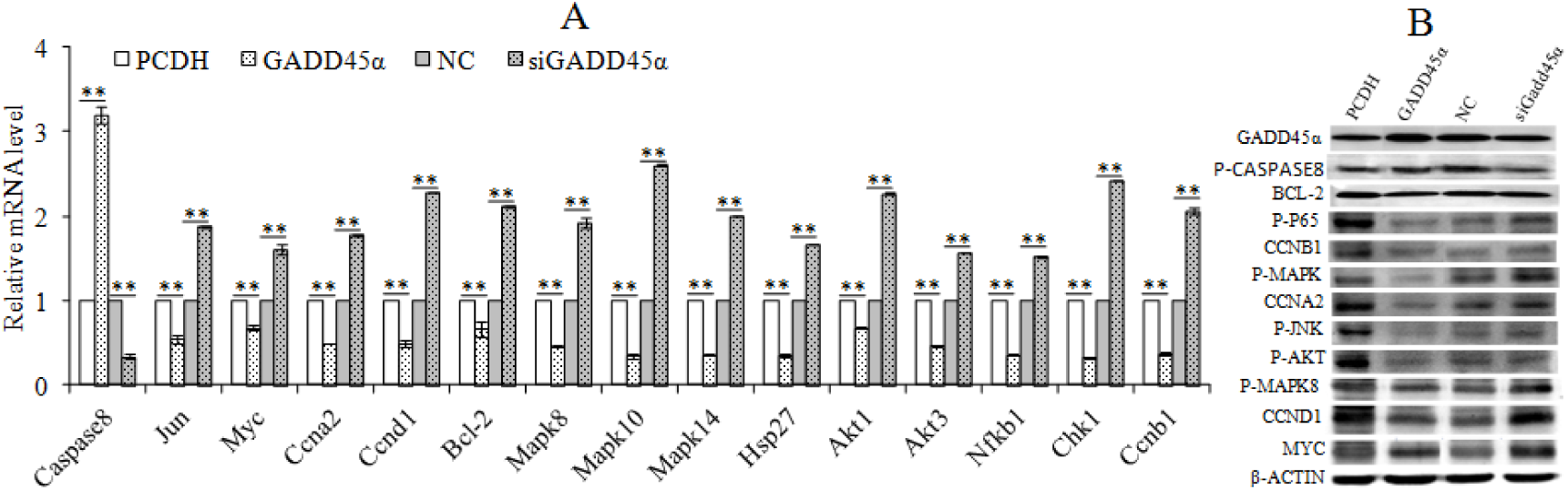
Expression changes of genes/proteins of GADD45α signaling pathway. A: genes expression level, B: proteins expression level * * *p*<0.01

The expression changes of genes-associated GADD45α signaling pathway were detected after treatment with 50μg/mL FZD or UVC for 30 s for 24 h. The results showed that overexpression of GADD45α could decreased the expression of Caspase3, Caspase8, Caspase9 and Bax at mRNA and protein in FZD and UVC-induced BRL-3A cells, and increased the expression of Myc, Bcl-2, Ccnd1, Ccnb1, PCNA and P21. While interfere with GADD45α expression increased the expression of Caspase8, Caspase9, Bax, Myc and Bcl-2 in FZD and UVC-induced BRL-3A cells, and decreased the expression of Ccnd1, Ccnb1, PCNA and P21 at mRNA and protein (Fig 15).

**Fig 15.**
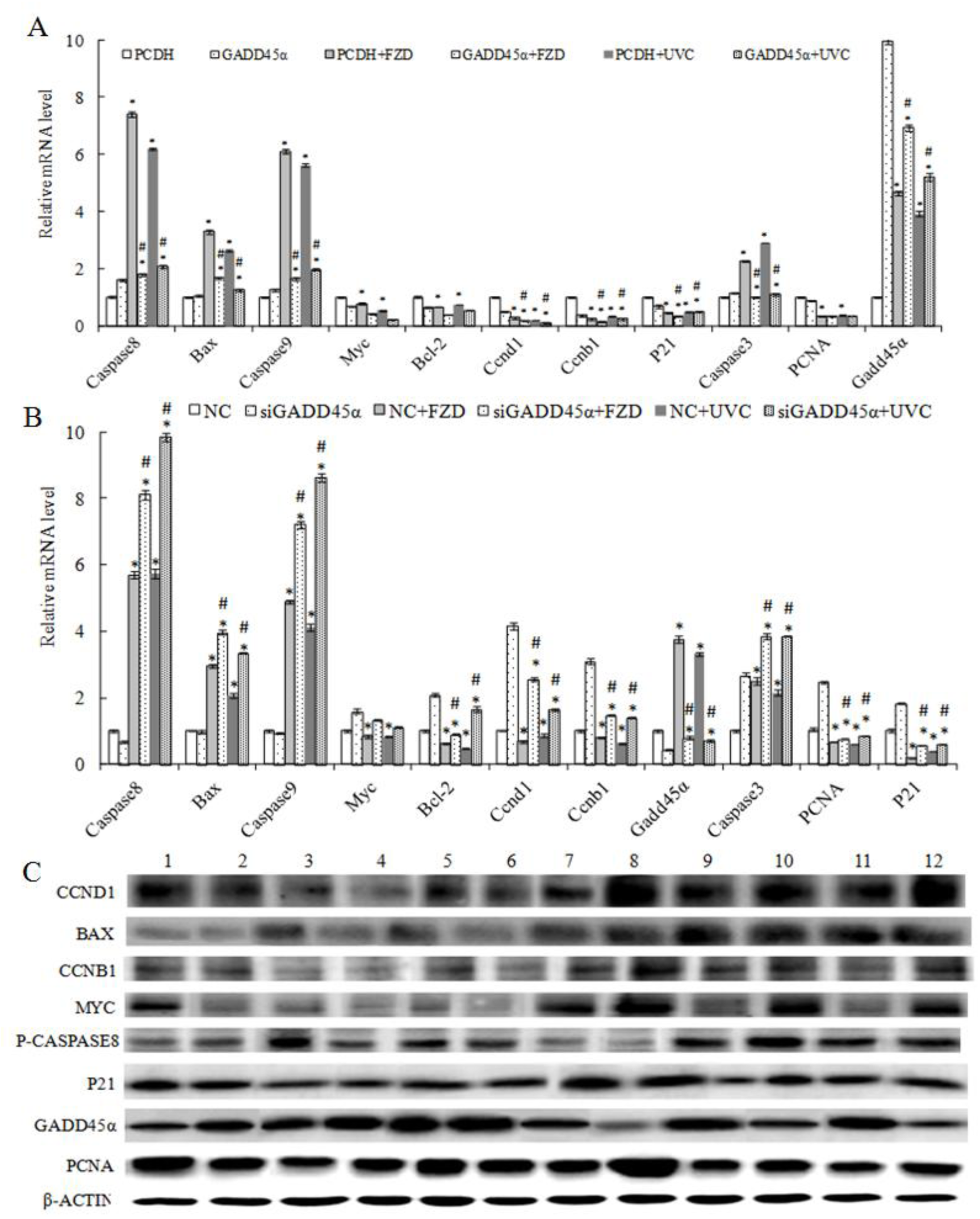
Expression changes of genes/proteins of GADD45α signaling pathway of BRL-3A cells that treated by FZD/UVC. A: Effect of up-regulated GADD45α expression on genes expression level; C: Effect of down-regulated GADD45α expression on genes expression level; C: proteins expression level. (* *p*<0.05 compared with control; # *p*<0.05 compared with control + FZD/UVC).

## 4. Discussion

Studies have shown that GADD45α was involved in cell cycle, apoptosis, genomic stability, DNA repair, immune response, and the occurrence of cancer. To understand the mechanism of GADD45α in liver regeneration, GADD45α signal pathway was constructed by several databases such as IPA, KEGG, QIAGEN and the related literature, gene addition and gene interference were used to investigate the effect of GADD45α on the cell viability, proliferation, apoptosis, cell cycle of BRL-3A and the expression of cell proliferation, apoptosis and GADD45α signal pathway-related genes/proteins.

In this paper, GADD45α up-regulated and down-regulated BRL-3A cells were obtained by lentivirus packaging and siRNA interference. The effects of GADD45α expression on the viability, proliferation, cycle and apoptosis of BRL-3A cells were detected by MTT, EdU and flow cytometry. The results showed that the viability, proliferation, that the number of cells in G1 and S phase, and the expression of proliferation-related genes Myc and Bcl2 of BRL-3A cells were decreased after GADD45α overexpression, while the number of G2/M phase and expression of apoptosisi-related genes Caspase8 were significantly increased. GADD45α down regulation could lead to the increase of cell viability, proliferation, the number of cells in G2/M and S phase, and Myc, Bcl2 expression, and the decrease of apoptosis rate and caspase8 expression. Our results indicated that GADD45α affects the activity and cell cycle of BRL-3A cells by proliferation, apoptosis-related genes/proteins, and ultimately regulates its proliferation and apoptosis.

In P38MAPK signaling pathway, GADD45α could inhibit P38MAPK via MKK3/6 (Yu et al., 2013), and regulated cell proliferation by MK2/3→ Iκb→NFκB (Dong et al., 2002; Tsai et al., 2003) or by Hsp27 (Su et al., 2008). Our study discovered that NFκB and Hsp27 were down-regulated at both mRNA and protein levels when overexpressing GADD45α, whereas NFκB and Hsp27 were up-regulated when GADD45α was interfered. Therefore, we believe that GADD45α may regulate BRL-3A cell proliferation through the P38MAPK pathway. In JNK pathway, GADD45α could inhibit JUN by JNK (David et al., 2002; Tront et al., 2006), and then regulate cell proliferation by FAS (Ivanov et al., 2001). The expression of MAPK8, JUN and FAS were down-regulated at mRNA and protein level after GADD45α up-regulated, and the expression of MAPK8, JUN and FAS were up-regulated at the mRNA level and protein level after GADD45α down-regulated, which indicated that GADD45α may regulate the proliferation of BRL-3A cells through JNK pathway. Studies have shown that in CDC2/CCNB1 pathway, GADD45α inhibited the kinase activity of the cdc2/cyclin B1 complex by interacting with cdc2 (Cdk1), which regulates the G2-M phase transition of the cell cycle (Liebermann and Hoffman, 2007). In this study, it was found that the expression of ccnb1 was down-regulated at both mRNA and protein levels when GADD45α was overexpressed, while interfered with the GADD45α expression, the expression of ccnb1 was upregulated, it was suggested that GADD45α may also regulate the cycle of BRL-3A cells through CDC2/CCNB1 pathway. Studies have shown that knockout of the GADD45α gene can reduce the expression of Akt protein in the lungs of mice (Meyer et al., 2009). The activated Akt could inhibit the activation of Bad and regulate apoptosis by Bcl-2 (Guo et al., 2011). GADD45α could inhibit the phosphorylation of STAT3 by interacting with mTOR, and inhibit tumor angiogenesis by blocking the mTOR/STAT3 pathway (Yang et al., 2013). The activation of STAT3 can induce the expression of the target gene c-myc, thereby promoting the expression of Cdc25a and promoting the process of cell cycle (Wang et al., 2011). In AKT and MTOR pathways, Akt, Bcl2 and Myc were down-regulated at mRNA level and protein level when GADD45α was overexpression. While GADD45α was down-regulated, the expression of Akt, Bcl2 and Myc in GADD45α expression was up-regulated, thus indicated that GADD45α may regulate the proliferation, apoptosis and cycle of BRL-3A cells by AKT and MTOR pathways. In order to investigate the mechanism of GADD45α on the proliferation and apoptosis of BRL-3A cells, the expression of GADD45α in BRL-3A cells was detected by qRT-PCR and Western Blot. The results showed that GADD45α regulate proliferation and apoptosis of BRL-3A cells by regulating the expression of Nfκb1, Mapk14, Mapk8, ccnb1, Akt1 and Myc, and by P38MAPK, JNK, CDC2/CCNB1, AKT and MTOR signaling pathways.

MTT, EdU and flow cytometry were used to detect the effect of GADD45α on cell viability, proliferation, cycle, DNA repair and apoptosis in FZD/UVC-induced BRL-3A cells. And the results indicated that GADD45α overexpression significantly increased the activity, cell proliferation, cell cycle arrest in S phase, and Myc, Bcl-2, Ccnd1, Ccnb1, PCNA, P21 expression of BRL-3A cells that treated by FZD/UVC, while decreased the apoptosis rate, and the expression of Caspase3, Caspase8, Caspase9 and Bax. In contrast, down-regulation of GADD45α expression by RNAi could reduce cell viability, cell proliferation, cell cycle arrest in S phase, and Myc, Bcl-2, Ccnd1, Ccnb1, PCNA, P21 expression in BRL-3A cells that treated by FZD/UVC, while increase cell apoptosis and the expression of Caspase3, Caspase8, Caspase9 and Bax.

GADD45α could regulate the expression of cell cycle regulatory genes (Kearsey et al., 1995), overexpression of GADD45α could induce cell cycle arrest (Maeda et al., 2002). DNA damage could enhance the activation of GADD45α, and induce cell cycle arrest and maintain genomic stability (Hollander et al., 1999). Gupta et al. found that knockout of the GADD45α gene increased the activation of Caspase-3 protein that induced by UVC radiation in mouse bone marrow cell. GADD45α and GADD45β interaction could promote cell survival by activating GADD45α-p38-NF-kappaB-mediated survival pathway and inhibiting GADD45β-mediated MKK4-JNK stress pathway in hematopoietic cells which exposed to UVC radiation (Gupta et al., 2006). FZD can effectively induce S-phase cell cycle arrest and inhibit cell growth, and increase oxidative induced DNA damage through ROS (Jin et al., 2011). These results indicate that GADD45α may play an important role in the activation of Myc, Bcl-2, Ccnd1, Ccnb1, PCNA and P21 in FZD/UVC-induced cell cycle arrest in BRL-3A cells. Sun et al. have shown that down-regulation of GADD45α expression in human liver tumor cells could reduce FZD-induced S-phase cell cycle arrest (Sun et al., 2015). Decreased GADD45 expression attenuated the inhibition of Cdc2/cyclinB1 activity in UV-irradiated human cells (Zhan et al., 1999). GADD45α inhibited Cdc2 kinase activity by altering cyclinB1 subcellular localization to induce cell cycle G2/M arrest and growth inhibition (Jin et al., 2002). Adenovirus-mediated expression of GADD45α is mediated by caspase activation and cell cycle arrest in G2/M phase (Li et al., 2009). Olaquindox treatment could increase the expression level of GADD45α protein and reactive oxygen species (ROS), and decrease the mitochondrial membrane potential (MMP), then increase the expression of Bax and decrease the expression of Bcl-2, and lead to the expression of cytochrome c (Cyt c). The down-regulation of GADD45α enhanced the production of ROS induced by quinoline indole, and then increases the activity of caspase-9, caspase-3 (Li et al., 2016).

In summary, GADD45α may regulate the proliferation, apoptosis and cycle of BRL-3A cells through P38MAPK, JNK, CDC2/CCNB1, AKT and MTOR signaling pathways. GADD45α may be involved in the DNA repair through Myc, Bcl-2, Ccnd1, Ccnb1, PCNA, P21, Caspase3, Caspase8, Caspase9 and Bax, and then regulate liver regeneration. This study is helpful to understand the mechanism of DNA damage repair that GADD45α regulated in liver regeneration process, thereby contributing to the prevention and treatment of liver disease. Further studies are still needed *in vivo* to confirm the mechanism of action of GADD45α.

## Acknowledgements

Funding: This work was funded by the National Natural Science Foundation Project of China (grant No. U1404312), the Science and technology project of Henan Province (grant No. 172102310507), the key scientific research project of universities in Henan Province (grant no. 13A180532), the doctoral Scientific Research Start-up Foundation of HNU (grant No. qd13033), and the Youth Science Foundation of HNU (grant no. 2014). I thank Associate Professor X.G. Yang for his suggestions and comments on the present manuscript.

## References

Bulavin, D. V., Kovalsky, O., Hollander, M. C. and Fornace, A. J., Jr. (2003). Loss of oncogenic H-ras-induced cell cycle arrest and p38 mitogen-activated protein kinase activation by disruption of Gadd45a. Mol Cell Biol 23, 3859–71.

Chang, Q., Bhatia, D., Zhang, Y., Meighan, T., Castranova, V., Shi, X. and Chen, F. (2007). Incorporation of an internal ribosome entry site-dependent mechanism in arsenic-induced GADD45 alpha expression. Cancer Res 67, 6146–54.

David, J. P., Sabapathy, K., Hoffmann, O., Idarraga, M. H. and Wagner, E. F. (2002). JNK1 modulates osteoclastogenesis through both c-Jun phosphorylation-dependent and -independent mechanisms. J Cell Sci 115, 4317–25.

Ding, Y., Chang, C., Niu, Z., Dai, K., Geng, X., Li, D., Guo, J. and Xu, C. (2016). Overexpression of transcription factor Foxa2 and Hnf1alpha induced rat bone mesenchymal stem cells into hepatocytes. Cytotechnology 68, 2037–47.

Dong, C., Davis, R. J. and Flavell, R. A. (2002). MAP kinases in the immune response. Annu Rev Immunol 20, 55–72.

Guo, Q., Jin, J., Yuan, J. X., Zeifman, A., Chen, J., Shen, B. and Huang, J. (2011). VEGF, Bcl-2 and Bad regulated by angiopoietin-1 in oleic acid induced acute lung injury. Biochem Biophys Res Commun 413, 630–6.

Gupta, M., Gupta, S. K., Hoffman, B. and Liebermann, D. A. (2006). Gadd45a and Gadd45b protect hematopoietic cells from UV-induced apoptosis via distinct signaling pathways, including p38 activation and JNK inhibition. J Biol Chem 281, 17552–8.

Hollander, M. C., Sheikh, M. S., Bulavin, D. V., Lundgren, K., Augeri-Henmueller, L., Shehee, R., Molinaro, T. A., Kim, K. E., Tolosa, E., Ashwell, J. D. et al. (1999). Genomic instability in Gadd45a-deficient mice. Nat Genet 23, 176–84.

Ivanov, V. N., Bhoumik, A., Krasilnikov, M., Raz, R., Owen-Schaub, L. B., Levy, D., Horvath, C. M. and Ronai, Z. (2001). Cooperation between STAT3 and c-jun suppresses Fas transcription. Mol Cell 7, 517–28.

Jin, S., Tong, T., Fan, W., Fan, F., Antinore, M. J., Zhu, X., Mazzacurati, L., Li, X., Petrik, K. L., Rajasekaran, B. et al. (2002). GADD45-induced cell cycle G2-M arrest associates with altered subcellular distribution of cyclin B1 and is independent of p38 kinase activity. Oncogene 21, 8696–704.

Jin, X., Tang, S., Chen, Q., Zou, J., Zhang, T., Liu, F., Zhang, S., Sun, C. and Xiao, X. (2011). Furazolidone induced oxidative DNA damage via up-regulating ROS that caused cell cycle arrest in human hepatoma G2 cells. Toxicol Lett 201, 205–12.

Kearsey, J. M., Coates, P. J., Prescott, A. R., Warbrick, E. and Hall, P. A. (1995). Gadd45 is a nuclear cell cycle regulated protein which interacts with p21Cip1. Oncogene 11, 1675–83.

Lapeyre, B., Bourbon, H. and Amalric, F. (1987). Nucleolin, the major nucleolar protein of growing eukaryotic cells: an unusual protein structure revealed by the nucleotide sequence. Proc Natl Acad Sci U S A 84, 1472–6.

Li, D., Dai, C., Zhou, Y., Yang, X., Zhao, K., Xiao, X. and Tang, S. (2016). Effect of GADD45a on olaquindox-induced apoptosis in human hepatoma G2 cells: Involvement of mitochondrial dysfunction. Environ Toxicol Pharmacol 46, 140–6.

Li, Y., Qian, H., Li, X., Wang, H., Yu, J., Liu, Y., Zhang, X., Liang, X., Fu, M., Zhan, Q. et al. (2009). Adenoviral-mediated gene transfer of Gadd45a results in suppression by inducing apoptosis and cell cycle arrest in pancreatic cancer cell. J Gene Med 11, 3–13.

Li, Z., Gu, T. P., Weber, A. R., Shen, J. Z., Li, B. Z., Xie, Z. G., Yin, R., Guo, F., Liu, X., Tang, F. et al. (2015). Gadd45a promotes DNA demethylation through TDG. Nucleic Acids Res 43, 3986–97.

Liebermann, D. A. and Hoffman, B. (2007). Gadd45 in the response of hematopoietic cells to genotoxic stress. Blood Cells Mol Dis 39, 329–35.

Maeda, T., Hanna, A. N., Sim, A. B., Chua, P. P., Chong, M. T. and Tron, V. A. (2002). GADD45 regulates G2/M arrest, DNA repair, and cell death in keratinocytes following ultraviolet exposure. J Invest Dermatol 119, 22–6.

Meyer, N. J., Huang, Y., Singleton, P. A., Sammani, S., Moitra, J., Evenoski, C. L., Husain, A. N., Mitra, S., Moreno-Vinasco, L., Jacobson, J. R. et al. (2009). GADD45a is a novel candidate gene in inflammatory lung injury via influences on Akt signaling. FASEB J 23, 1325–37.

Michalopoulos, G. K. and DeFrances, M. C. (1997). Liver regeneration. Science 276, 60–6.

Saintigny, Y., Delacote, F., Vares, G., Petitot, F., Lambert, S., Averbeck, D. and Lopez, B. S. (2001). Characterization of homologous recombination induced by replication inhibition in mammalian cells. EMBO J 20, 3861–70.

Su, X., Ao, L., Zou, N., Song, Y., Yang, X., Cai, G. Y., Fullerton, D. A. and Meng, X. (2008). Post-transcriptional regulation of TNF-induced expression of ICAM-1 and IL-8 in human lung microvascular endothelial cells: an obligatory role for the p38 MAPK-MK2 pathway dissociated with HSP27. Biochim Biophys Acta 1783, 1623–31.

Sun, Y., Tang, S. and Xiao, X. (2015). The Effect of GADD45a on Furazolidone-Induced S-Phase Cell-Cycle Arrest in Human Hepatoma G2 Cells. J Biochem Mol Toxicol.

Tront, J. S., Hoffman, B. and Liebermann, D. A. (2006). Gadd45a suppresses Ras-driven mammary tumorigenesis by activation of c-Jun NH2-terminal kinase and p38 stress signaling resulting in apoptosis and senescence. Cancer Res 66, 8448–54.

Tsai, P. W., Shiah, S. G., Lin, M. T., Wu, C. W. and Kuo, M. L. (2003). Up-regulation of vascular endothelial growth factor C in breast cancer cells by heregulin-beta 1. A critical role of p38/nuclear factor-kappa B signaling pathway. J Biol Chem 278, 5750–9.

Vairapandi, M., Azam, N., Balliet, A. G., Hoffman, B. and Liebermann, D. A. (2000). Characterization of MyD118, Gadd45, and proliferating cell nuclear antigen (PCNA) interacting domains. PCNA impedes MyD118 AND Gadd45-mediated negative growth control. J Biol Chem 275, 16810–9.

Vairapandi, M., Balliet, A. G., Hoffman, B. and Liebermann, D. A. (2002). GADD45b and GADD45g are cdc2/cyclinB1 kinase inhibitors with a role in S and G2/M cell cycle checkpoints induced by genotoxic stress. J Cell Physiol 192, 327–38.

Wang, H., Lafdil, F., Kong, X. and Gao, B. (2011). Signal transducer and activator of transcription 3 in liver diseases: a novel therapeutic target. Int J Biol Sci 7, 536–50.

Wang, X. W., Zhan, Q., Coursen, J. D., Khan, M. A., Kontny, H. U., Yu, L., Hollander, M. C., O’Connor, P. M., Fornace, A. J., Jr. and Harris, C. C. (1999). GADD45 induction of a G2/M cell cycle checkpoint. Proc Natl Acad Sci U S A 96, 3706–11.

Yang, F., Zhang, W., Li, D. and Zhan, Q. (2013). Gadd45a suppresses tumor angiogenesis via inhibition of the mTOR/STAT3 protein pathway. J Biol Chem 288, 6552–60.

Yu, Y., Li, J., Wan, Y., Lu, J., Gao, J. and Huang, C. (2013). GADD45alpha induction by nickel negatively regulates JNKs/p38 activation via promoting PP2Calpha expression. PLoS One 8, e57185.

Zhan, Q., Antinore, M. J., Wang, X. W., Carrier, F., Smith, M. L., Harris, C. C. and Fornace, A. J., Jr. (1999). Association with Cdc2 and inhibition of Cdc2/Cyclin B1 kinase activity by the p53-regulated protein Gadd45. Oncogene 18, 2892–900.

